# Rare Microbial Taxa Emerge When Communities Collide: Freshwater and Marine Microbiome Responses to Experimental Seawater Intrusion

**DOI:** 10.1101/550756

**Authors:** Jennifer D. Rocca, Marie Simonin, Justin P. Wright, Alex Washburne, Emily Bernhardt

## Abstract

Whole microbial communities regularly merge with one another, often in tandem with their environments, in a process called community coalescence. Such events allow us to address a central question in ecology – what processes shape community assembly. We used a reciprocal transplant and mixing experiment to directly and independently unravel the effects of environmental filtering and biotic interactions on microbiome success when freshwater and marine communities coalesce. The brackish treatment and community mixing resulted in strong convergence of microbiome structure and function toward the marine. Brackish exposure imposed a 96% taxa loss from freshwater and 66% loss from marine microbiomes, which was somewhat counterbalanced by the emergence of tolerant rare taxa. Community mixing further resulted in 29% and 49% loss from biotic interactions between freshwater and marine microbiomes, offset somewhat by mutualistically-assisted rare microbial taxa. Our study emphasizes the importance of the rare biosphere as a critical component of community resilience.

## INTRODUCTION

A fundamental goal in ecology is to determine the distribution and abundance of species and the mechanisms controlling this distribution. This central objective is particularly challenging in microbial ecology because of the immense diversity, brief lifespan, and microscopic size of microorganisms. Many studies examine organismal distributions along environmental gradients to shed light on the natural history of species (Whittaker 1965) and microbial ecologists also use this concept to understand the natural history of microorganisms at biogeographical scales (Fierer and Jackson 2006) and along elevational (Bryant *et al.* 2008; Yang *et al.* 2014; Siles and Margesin 2016), precipitation (Angel *et al.* 2009; Hawkes *et al.* 2017) and salinity (Herlemann *et al.* 2011; Herlemann *et al.* 2016) gradients. However, distinguishing between environmental tolerance or competitive abilities for microorganisms and determining their fundamental niche is difficult to assess with just distribution information. Determining the relative importance of environmental filtering and biotic interactions in structuring extant communities is difficult as both processes operate concurrently (Vellend 2010; Goberna *et al.* 2014; Cadotte and Tucker 2017).

Quantifying the influence of these assembly filters is unique for microbial communities, which typically migrate in the aggregate and in tandem with their environment. Thus ‘habitat patches’ move along with an entire assemblage of organisms. In metacommunity ecology, such collective exchanges are referred to as mass effects (Leibold *et al.* 2004, Souffreau *et al.* 2014, Comte *et al.* 2017). When previously distinct communities combine along with their respective environments, the reassembly of the novel community is termed ‘community coalescence’ (Webb 1976, Livingston *et al.* 2013, Rillig *et al.* 2015, Rillig & Mansour 2017). Microbial community coalescence occurs every time a leaf falls to the ground, a soil particle is blown into a new landscape, or two bodies of water mix. Despite the ubiquity of microbial community coalescence, the formal recognition of this concept is fairly recent in microbial ecology (Rillig *et al.* 2017; Mansour *et al.* 2018). In fact, Mansour *et al.* (2018) suggest that many experimental studies with microbial communities are unacknowledged community coalescence experiments.

Community coalescence is not only an interesting phenomenon, it also provides immense opportunities to leverage experiments to learn about <u>*how*</u> individual microorganisms respond and microbial communities assemble in response to the merging event. To develop the natural history of microorganisms, we need to understand how both environmental and biological filters ultimately determine who persists and who does not when communities collide. To address this fundamental ecological question, we introduce a new methodological framework (Fig. 1) to directly measure the independent and combined effects of environmental and biotic filters (sensu Vellend 2010) that structure the distribution and abundance of microbial taxa.

**Figure 1.**
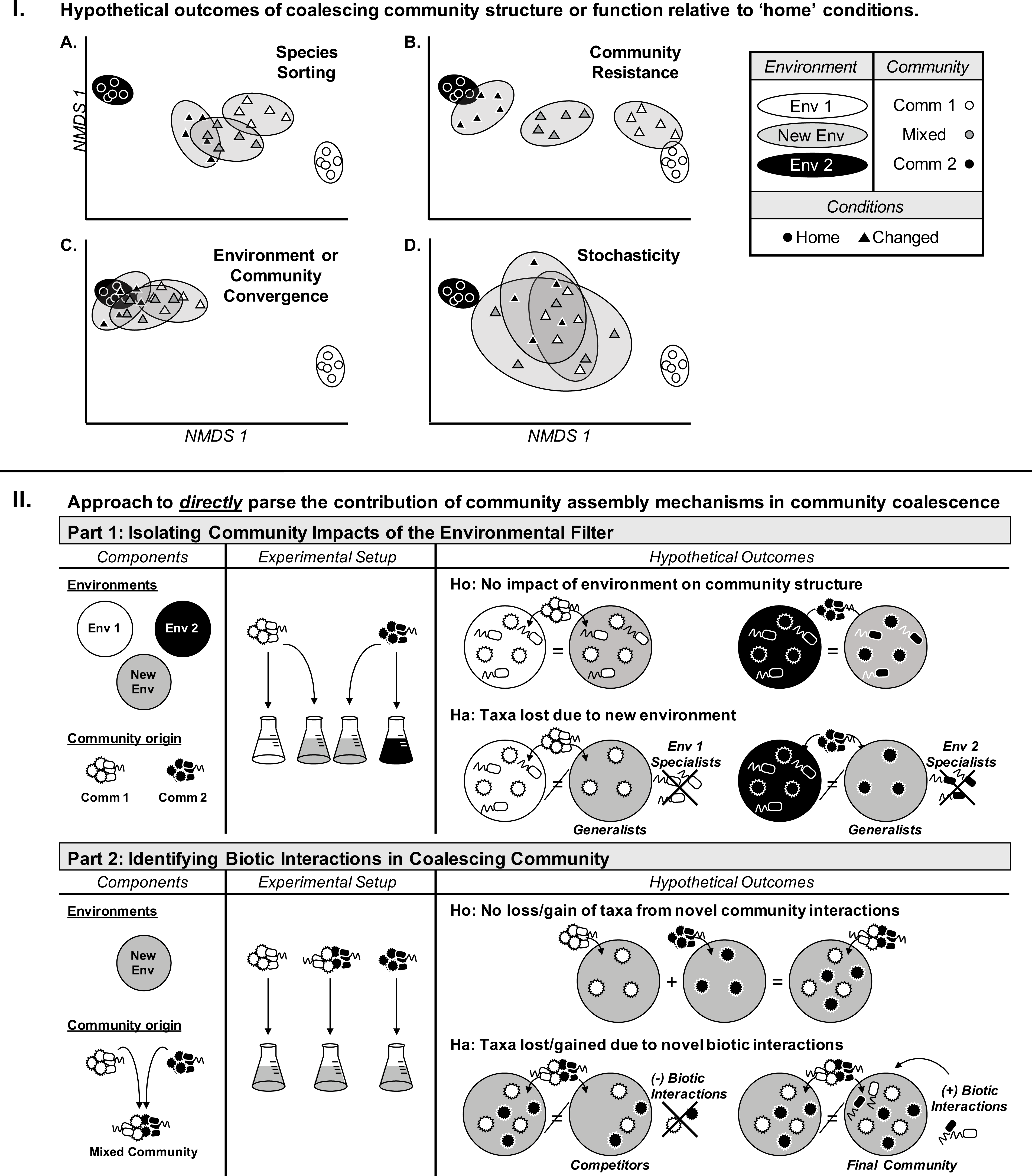
**(I.) Potential outcomes of coalescing communities relative to “home” conditions.** Ellipsoids represent the theoretical variation in community structure or function among experimental replicates (symbols). Communities may be symmetrically impacted by coalescence (A,B), constrained primarily by environment (A), or resistant to environmental change (B); (C) asymmetrical impact may be driven by either environment or community; or (D) coalescence may result in stochastic shifts in community structure or function. **(II.) Conceptual framework to disentangle the influence of assembly processes**: components needed for each filter (left), experimental setup (center), and hypothetical outcomes (right). Top panel: assessing the impact of the environmental filter, where no impact of altered environment (Ho), versus a shift in community structure (Ha), reveals environmentally sensitive and generalist taxa. Bottom panel determines the impact of novel biotic interactions, with no impact (Ho) where the sum of Part 1 communities equals that of the coalesced community, versus new biotic interactions, where taxa are lost due to competition or gained due to mutualistic interactions (Ha).

We used a microcosm experiment to examine how microbial communities reassemble following the blending of media and microbial communities between freshwater and marine environments to simulate seawater intrusion - a mixing event that occurs along all coastal margins. Our experimentally created coalescence event (Fig. 1) allowed us to ask: Q1) how do microbial communities from distinct environments respond to merging? Q2) what is the relative role of environmental filtering and biotic interactions in structuring this newly united community? Q3) are more closely related taxa more likely to respond similarly to each filter? and Q4) what are the resulting consequences for microbial community function?

Seawater intrusion into freshwater systems is a prevalent feature of tidal environments and is increasing in frequency and extent as drought, irrigation and climate change are all increasing the inland and upland movement of seawater into freshwater habitats (Weston *et al.* 2006, Herbert *et al.* 2015). The multivariate chemical transition to brackish water is well studied, and salinity is a well-documented environmental stressor (Lozupone and Knight 2007), imposing strong evolutionary selection on organisms (Paver *et al.* 2018). When freshwaters come into contact and blend with seawater, we observe dramatic decreases in organic carbon content due to complexation with salts, increased pH, and nutrient availability along with increases in salinity (Craft *et al.* 2009, Barlow & Reichard 2010, Ardón *et al.* 2016a). Dynamic pulses of salty, oligotrophic marine water mixing with relatively mesotrophic freshwater habitats results in brackish conditions, novel to either endpoint microbial community. The microbial consortia derived from either endpoint habitat are well adapted to their respective environmental conditions (Canfora *et al.* 2014, Herlemann *et al.* 2016), and exposure to brackish water imposes substantial stress on members of each endpoint community, particularly the freshwater sediment microbial communities (Baldwin *et al.* 2006, Jackson and Vallaire 2009, Neubauer *et al.* 2013,) and water column microbial taxa (Burke and Baird 1931, Nielsen *et al.* 2003, Ewert and Deming 2013). While we expect freshwater microorganisms to more stressed by the salt exposure of the brackish conditions (Edmonds *et al.* 2009, Herbert *et al.* 2015), little is known about the outcome of blending these distinct communities. We expect a substantial turnover of each microbiome in response to the strong environmental filter imposed by the brackish exposure (Paver *et al*. 2018), but less is known about the community responses to novel biotic interactions that might arise through community blending. Though marine microbial taxa are more stress tolerant to the wide range of salinity, they may not fare well if the freshwater microbial taxa are better competitors (Grimes *et al.* 1977). Do the existing communities have a great phenotypic plasticity or can the rare biosphere aid in the resilience to drastic environmental changes and community introductions?

Our experimental design allows us to differentiate amongst four potential outcomes of microbial community coalescence (Fig. 1, top panel). Beginning with the assumption that our starting endmember communities are distinct, we expect that the community resulting from their exposure to the mixed environment and to the community merger would follow one of four possible trajectories of the final community composition (Fig. 1, top panel): A) intermediates between the two home conditions due to species sorting hinging on the environment, B) stratified yet similar to their initial inocula due to symmetric community resistance; or C) converge towards the assemblage observed in only one of the two environments due to strong asymmetric environmental selection <u>*or*</u> asymmetric community resistance. Alternatively, a final outcome (D) is that the coalescence conditions result in highly variable emergent communities via stochastic responses. In the scenario where the communities converge towards a single endmember (C), our methodological framework allows us to directly distinguish between environmental and biotic controls in driving this response (Fig. 1, bottom panel).

Our integrative research approach allows us the opportunity to add a natural history perspective to the study of microbial ecology. Habitat transplants will apply an environmental filter to identify generalist vs specialist microbial taxa. Community coalescence allows us to identify strong vs. weak competitors and common associations amongst taxa.

## MATERIALS and METHODS

### Field sample collection and aquatic endmember characterization

The two endmember sources for this microcosm experiment are located in coastal North Carolina, USA (Table 1). The Freshwater Wetland site, hereafter “Freshwater” site, is a blackwater wetland ecosystem located within the Timberlake Wetland Restoration Project in Tyrrell Co., NC, 35°53’46.4"N, 76°09′51.4″W (Ardón *et al.* 2016b), exposed to episodic or storm-triggered flows following rain or coastal storm systems. In contrast, the Coastal Marine site, hereafter “Marine” site, located at the northern end of the Cape Hatteras National Seashore (Dare Co., NC, 35°49’57.4"N, 75°33′25.7″W) with persistent water turbulence.

The Freshwater and Marine endmember sites were selected for their close proximity (64 km) and even latitude, while maximizing the potential range of habitats involved in seawater intrusion. Both sites are not significantly impacted by human activity, and the Freshwater site had no recent seawater exposure. Additionally, the water samples were specifically collected at the end of a seasonal period where recent seawater intrusion was least likely. We acknowledge that these sites do not represent true potential locations of seawater intrusion, as the mixing occurs gradually across the 64 km gradient. However, the sites are true endmembers, with past seawater intrusion coalescence events. In June 2016, we collected 80 L of surface water from each site into sterile carboys, stored at the average temperature (23 ^o^C) of the two sites at time of collection. Within 12 hours, each sample was filtered through axenic 1 mm mesh to remove macroorganisms and debris. Each sample was subsampled for salinity, pH, and dissolved organic carbon. After filtration, the two endmember water samples were halved to generate: (a) the microbe-free environments and (b) microbial inocula (Fig. 1).

**Table 1.** Location, chemical and microbial bulk characterization of the two distinct aquatic endmember habitats in Coastal North Carolina, USA: Freshwater and Marine environments and inocula; and the chemical characterization of the experimentally mixed (1:1) Brackish environment and blended inoculum.

| *Environment* | *Freshwater* | *Brackish* | *Marine* |
| --- | --- | --- | --- |
| *Endpoint Location* | Timberlake Restoration | Lab mixed 50:50 by volume | Cape Hatteras NS |
| Latitude, | 35°53'46.4"N, | — | 35°49'57.4"N, |
| Longitude | 76°09'51.4"W | — | 75°33'25.7"W |
| Mixing | Episodic/Storm | — | Turbulent |
| Temp (June '16) | 23-25 °C | — | 23 °C |
| *Chemistry of Environments* |  |  |  |
| pH | 4.4 (± 0.05) | 7.8 (± 0.01) | 8.2 (± 0.01) |
| Salinity (PSU) | 0.07 (± 0.01) | 26.7 (± 0.9) | 51.8 (± 0.4) |
| Dissolved Organic Carbon (mg/L) | 33.9 (± 1.2) | 21.7 (± 3.2) | 3.6 (± 0.4) |
| Dissolved Inorganic Nitrogen (mg/L) | 0.92 (± 0.01) | 0.61 (± 0.05) | 0.12 (± 0.03) |
| *Microbial Inoculum Characterization* |  |  |  |
| Biomass (ng DNA/μL) | 749.7 | 584.8 | 218.3 |

### Microcosm incubation setup

We set up a laboratory incubation, exposing three inocula isolated from each endmember samples (Freshwater, Marine, and a 1:1 mixture of Freshwater-Marine, hereafter “Coalescence”), into three axenic aquatic environments (Freshwater, Marine, and a 1:1 mixture of the endmember environments, hereafter “Brackish”). We sterilized each water source by autoclaving at 121 ^o^C / 20 PSI for 30 mins in covered acid-washed glassware. After cooling to room temperature, the autoclaving was repeated twice more to ensure that any microorganisms exiting dormancy were killed. Subsamples of each starting environment were streaked on Luria-Bertani agar plates to confirm sterility.

Intact microbial communities were isolated by concentrating cells off the remainder of the Freshwater and Marine samples. Two complementary concentrating methods were implemented to minimize biases imposed by either method. Half from each environment was gently centrifuged at 5,000 RCF in 50 mL batches in round-bottom tubes to ensure maximal viability of the microbial assemblages (Pembrey *et al.* 1999, Peterson *et al.* 2012). In parallel, the remaining water samples were filtered over gamma-irradiated Pall Supor 0.2 µm nitrocellulose membranes (Millipore, New York, NY) in small batches to minimize fouling on the filter. For each inoculum, the filter-collected and centrifuged-collected microbiomes were combined by resuspending in sterilized home. The Coalescence microbiome was made by mixing 1:1 subsamples of the Freshwater and Marine inocula.

The full experimental design consisted of 120 independent microcosms (850 mL inoculated with 550 µl), with an experimental replication of five, along with positive/negative controls (Suppl. Fig. 1). Here we present a subset (n=25 microcosms) of the full incubation with five treatment conditions: “Freshwater-Home”: Freshwater microbiome in Freshwater environment, “Marine-Home”: Marine microbiome in Marine environment, “Freshwater-Brackish” and “Marine-Brackish”: Freshwater (or Marine) microbiome in Brackish environment; and “Brackish-Coalescence”: a 1:1 blended microbiome in a 1:1 mixture of the endmember environments. We set up the microcosms into sterile glass Mason jars under UV-treated PCR-hood conditions. The positive control microcosms were used to correct for any experimental artifacts from inoculum and environment preparations, which did not vary significantly from the ‘home’ microcosms (Suppl. Fig. 2). Microcosms with sterile environments or sterile water with no added inocula were included as negative controls to confirm axenic conditions, and none had any detectable microbial growth or genetic material and will not be discussed further. The incubation ran for seven days in an environmental growth chamber (23 ^o^C, 13.5 hr diurnal light regime, and PAR: 250-450 µmols m^-2^s^-1^) reflecting field conditions. Twice-daily, the microcosms were re-randomized to minimize potential biases from differential light and temperature across the chamber, and each microcosm was mixed by inverting.

**Figure 2.**
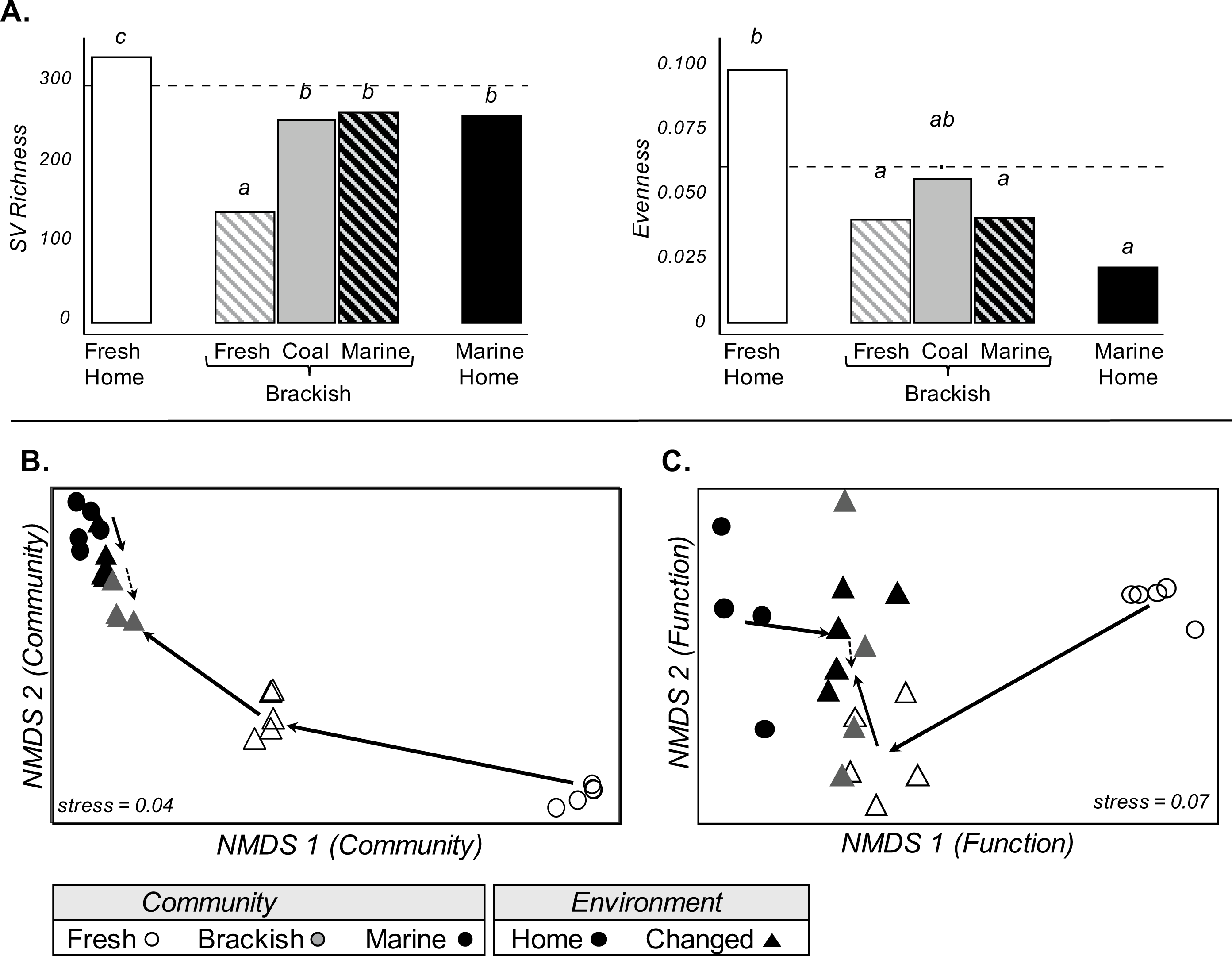
Impact of coalescence on microbial community structure and function: (A) alpha diversity of each treatment for SV richness (left) and Pielou’s Evenness (right), (B,C) nonmetric multidimensional scaling (NMDS) of (B) bacterial community structure; and (C) microbial extracellular enzyme activity. The NMDS arrows indicate movement through the filters relative to endmember control conditions, weighted by significance from Adonis (solid = significant change; dotted = NS). No community or functional data is available for one Brackish-Coalescent sample that was damaged during the incubation (n=4, instead of experiment wide n=5).

### Microbial function

Microbial extracellular enzyme potential activity (hereafter enzyme activity) was measured in each microcosm at the end of the incubation. Enzyme activity for eight enzymes were measured from a 91-mL homogenized subsample of each microcosm following a protocol developed by Bell *et al.* (2013), here modified to handle water. The following enzymes were targeted with fluorescently labeled substrates to capture potential N, P, C, and S degradation activity: *α*-1,4-glucosidase (AG), aryl-sulfatase (AS), *β*-1,4-glucosidase (BG), *β*-D-1,4-cellobiosidase (CB), L-leucine aminopeptidase (LAP), *β*-1,4-N-acetylglucosaminidase (NAG), alkaline phosphatase (PHOS), and *β*-D-xylosidase (XYL). After three-hours at room temperature, the centrifuged supernatant for each sample was read (340/460 nm) on a FLUOstar Optima spectrophotometer (BMG Labtech, Cary, NC, USA) in black optical 96-well plates.

### DNA extraction & bacterial community analysis

After the incubation, a 250 mL subsample of each microcosm was filtered over gamma-irradiated Pall Supor 0.2 µm nitrocellulose membranes (Millipore, New York, NY), and the filtrate was centrifuged at 10,000 x g for one hour to pellet any remaining ultra-small microorganisms (Luef *et al.* 2015). The pellet and filter were combined and processed for genomic DNA using a MoBio PowerWater DNA Isolate Kit (MoBio, Vancouver, CA) modified by adding a heating step during cell lysis. Total genomic DNA was fluorometrically measured (Quant-iT dsDNA Assay Kit, ThermoScientific, Waltham, MA), and used as proxy for total microbial biomass (Baas *et al.* 2015, Nagler *et al.* 2018). The samples were amplified, targeting the V4 hypervariable region of the bacterial 16S rRNA gene (515-F/806-R, Caporaso *et al.* 2011), and sequenced with Illumina MiSeq (PE 150bp; V2 chemistry) at the Environmental Sample Preparation and Sequencing Facility (ESPSF) at Argonne National Laboratory. Raw sequences are deposited into the NCBI Sequence Read Archive (SRA): PRJNAXXXX.

ESPSF returned 25 million raw sequences, which we processed through Quantitative Insights Into Microbial Ecology 2 (Qiime2) pipeline (Bolyen *et al.* 2018) to remove low quality reads and putative chimera, and to denoise the sequences into exact sequence variants (SVs) with Dada2 (Callahan *et al.* 2017). We aligned the representative sequences and assigned taxonomy using the Silva V132 (99%) curated reference alignment (Quast *et al.* 2013), and a phylogeny and improved alignment were simultaneously generated using the Practical Alignment using SATé and TrAnsitivity (PASTA) software (Mirarab *et al.* 2013). Eukaryotic, mitochondrial, and chloroplast contaminant sequences were removed. The SV table was rarefied to lowest sequence depth (17,500 sequences), and the final dataset contained 3389 unique SVs with 12,293,437 total reads.

### Microbiome Structure & Function Profile

To bulk characterize the bacterial communities, alpha-diversity (Chao1) and Pielou’s evenness were calculated on each sample. A Bray-Curtis dissimilarity matrix was created for the community dataset (and Euclidean for the enzyme dataset) to examine differences in community structure and in the functional profile among treatments, and to visualize these shifts using non-metric multidimensional scaling (NMDS). To test the hypothetical outcomes of community structure detailed in Fig. 1A, a non-parametric multivariate analysis of variance (per-MANOVA, Anderson 2001) was implemented to identify significant overall and between group shifts in community. The “adonis” function in the vegan R package (Version 3.3.1, Oksanen *et al.* 2017) was used to implement the perMANOVA by estimating correlation coefficients and corresponding p-values (permutations = 999) for the effect of each treatment. Subsequent pairwise comparisons were performed using Adonis. The “procrustes" function (Jackson 1995) in Vegan was used to perform least-squares orthogonal mapping to determine correlations between two multivariate datasets. Here, procrustes (PROTEST) was used to determine correlations between bacterial community structure and microbial enzyme activities profile. Additionally, we visualized the responses of the most abundant microbial SVs (> 2%) using UPGMA clustering (Sokal & Michener 1958) to the environmental and microbiome mixing treatments. This subset of taxa was further identified for halotolerance using the LPSN database (Parte 2013) of known microbiological traits. Univariate data that violated assumptions of normal distribution and homoscedasticity were log-transformed, and assumptions were subsequently re-verified with Shapiro-Wilk and Levene’s tests and examination of Q-Q plots. We used one-way ANOVAs to test the significance of the treatments on microbiome diversity and enzyme activity, then *post-hoc* pairwise comparisons among the treatment combinations were performed using Tukey’s Honestly Significant Difference multiple means comparison.

### Direct separation of contribution of local assembly mechanisms

To directly disentangle the influence of environmental filtering and novel biotic interactions, we compared specific experimental treatments (Fig. 1, bottom panel). To examine the environmental filter (<u>*Part 1*</u>), we compared ‘home’ microbiomes to the corresponding Brackish microbiomes (i.e. Freshwater-Home vs. Freshwater-Brackish or Marine-Home vs Marine-Brackish). To examine the impact of novel biotic interactions from community blending (<u>*Part 2*</u>), the Freshwater-Brackish and Marine-Brackish microbiomes were compared to the Brackish-Coalescence microbiomes. A new phylogenetic-based method – phylofactorization – was used to identify clades driving changes in community composition (Washburne *et al* 2017, 2019), based on a Holm’s sequentially rejective 5% cutoff for the family-wise error rate. The resultant phylofactor objects are available in the supplementary material.

## RESULTS

### Starting Conditions: Bacterial Community and Chemical Characterization

Our two environments and initial microbial communities were distinct among the Freshwater and Marine endmembers (Table 1). Other than water temperature at the time of field collection, all other water chemistry properties varied substantially between our endmembers (Table 1). Electrical conductivity was ~740-fold greater and pH was 3.8 units higher in the Marine sample, while the Freshwater environment had higher concentrations of both nutrients and dissolved organic matter (Table 1).

The Freshwater and Marine microbial communities were also distinct from one another. Microbial biomass (Table 1: 749 vs. 218 ng DNA/µL), alpha diversity (Fig. 2A: 328 vs. 253 observed SVs/microcosm) and evenness (Fig. 2A: 0.1 vs. 0.02) were significantly higher in the Freshwater microbial community. The two endmember microbiomes had minimal overlap in their community composition (Fig. 2B), with only 21 SVs (<1.4% of total SVs) representing 22% of the Fresh+Marine summed biomass (Suppl. Fig. 3). The Freshwater community was dominated by the families: *Acetobacteraceae, Paracaedibacteraceae, Beijerinckiaceae, Burkholderiaceae*; while the Marine microbiome was dominated by the families: *Alteromonadaceae, Rhodobacteraceae, Saprospiraceae, Spirosomaceae*, and *Vibrionaceae* (Fig. 3) with a single taxon – *Alteromonas* - dominating 9.7% of the Marine microbiome. The two communities contrasted in their functional potential as well, exhibiting distinct enzyme profiles, with the Freshwater microbiome producing significantly higher amounts in seven of the eight enzymes analyzed (Fig. 2C, Suppl. Fig. 4).

**Figure 3.**
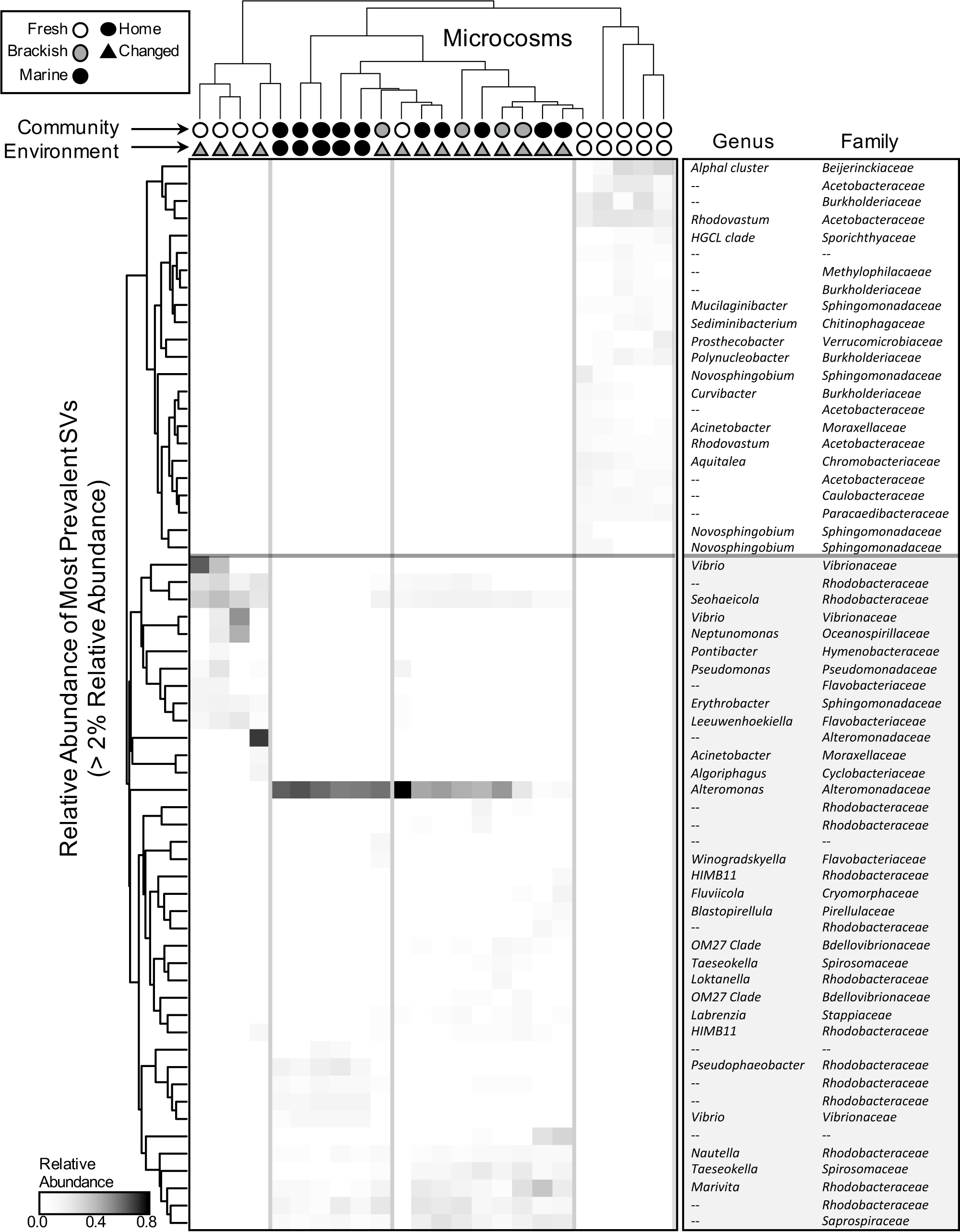
Responses of the most abundant bacteria to community coalescence. Heatmap of the most prevalent bacterial SVs (>0.5% relative abundance among all microcosms), vertically clustered by SV, with corresponding taxonomy (shaded by salt tolerance) and horizontally by microcosm, identified by Environment (top row symbols) and Inoculum (bottom row). Blue and green flanked regions identify the “home” conditions for Freshwater and Marine conditions. Ranking estimated using standard UPGMA methods based on Bray-Curtis pairwise dissimilarities in Vegan R package.

**Figure 4.**
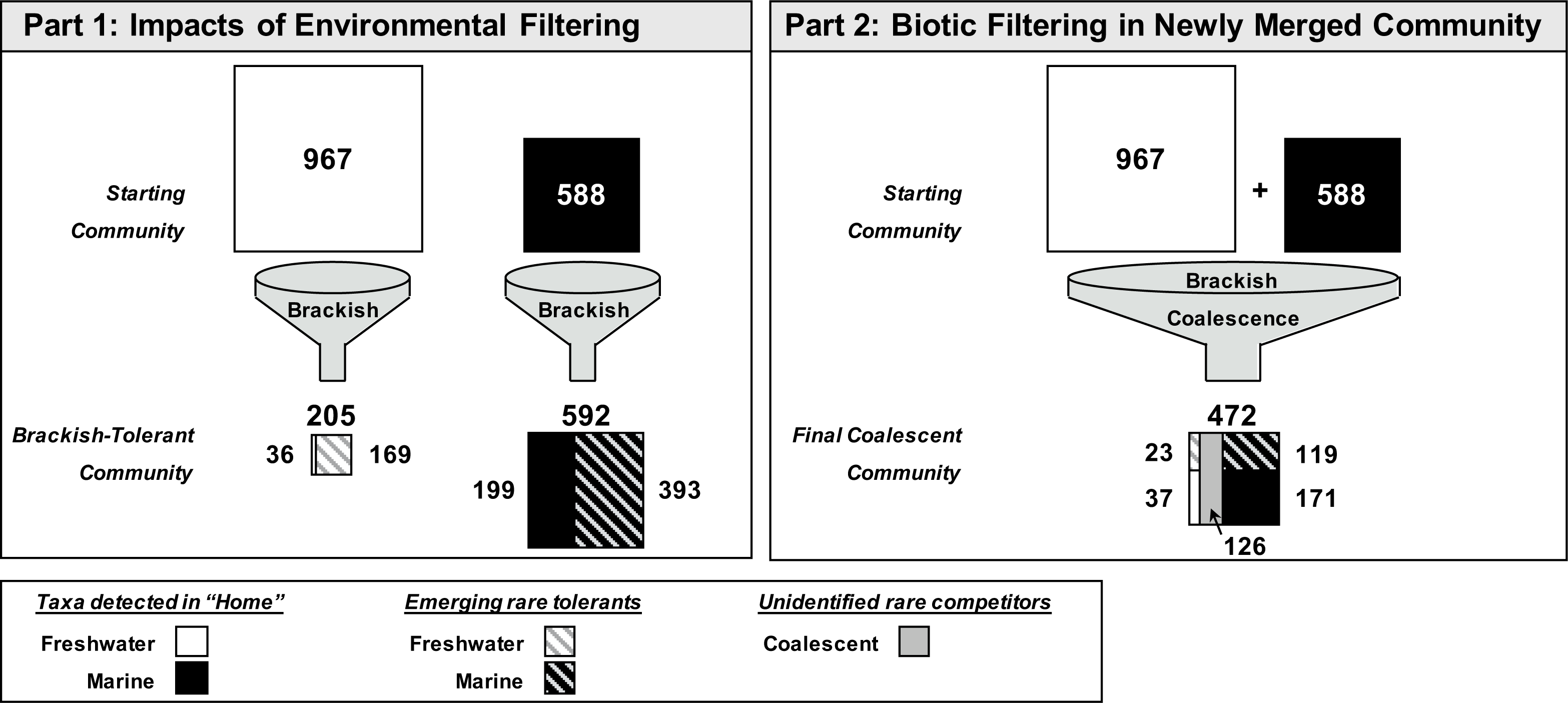
Distribution of community origin due to each assembly filter: microbial taxa lost and gained due to Brackish exposure (Part 1: Impacts of Environmental Filtering); and due to microbiome merging with the Brackish-Coalescence treatments (Part 2: Biotic Filtering In Newly Merged Community). The numbers inside or above each box represent the number of taxa (SVs) found among the microcosms of that treatment (n=5), with Freshwater in “white” and Marine in “Black” boxes. The numbers adjacent to the lower boxes represent the number of taxa detected in the original communities (solid) or detected as rare taxa emergence (striped), and the gray box represents rare emergence of unknown endmember origin.

### Convergence towards Marine Bacterial Community During Coalescence

When the two axenic endmember water samples were combined to create our Brackish media, the resulting chemical properties were intermediate to the two endmembers. Substantial buffering led to the Brackish media having a pH closer to the endmember, but other chemical components, were essentially the average of the two contributing media (Table 1). The blending of the two communities into the Coalescence inoculum was also the average biomass of the two endmembers (Table 1).

*Environmental Filter:* Despite their higher diversity and biomass (Fig. 2A), the Freshwater microbial taxa did not fare well when added to the Brackish media in the absence of Marine community blending (i.e. Freshwater-Brackish treatment). Only 36 of the 967 total taxa initially sequenced from our Freshwater-Home microbiomes persisted following this environmental filter into Brackish media, although 169 taxa that were below our detection (i.e. rare taxa) in the initial inoculum were detected in the Brackish media (Fig. 4). We are confident that these taxa represent increases in abundance from the rare biosphere contained in the initial inoculum as we failed to detect any genomic DNA in our negative controls. Marine microbial taxa were more tolerant of the transfer into Brackish media, with 199 of the original 588 taxa surviving. A large number of rare biosphere taxa emerged from Marine inoculum under Brackish conditions. We detected 393 taxa in our Marine-Brackish treatments that were not detected in the original Marine-Home microcosms. Consequently, the diversity of the Marine inoculum in Brackish media was equal to the starting inoculum (592 vs. 588), while there was a substantial loss of total richness for the Freshwater microbiome in Brackish media compared to the diversity of its initial inocula (205 vs 967) (Fig. 4). The community composition of the Marine-Brackish replicates was very similar to the Marine-Home, while the Freshwater-Brackish community shifted significantly in composition towards the Marine-Home relative to its initial Freshwater-Home composition (Fig. 2B). We detected more shared taxa between these environmentally filtered communities, with 65 taxa overlapping between Freshwater-Brackish and Marine-Brackish. For the low diversity Freshwater-Brackish replicates, these shared taxa represent more than 25% of the total diversity. These increased taxa include the following families, which were below detection limit in Freshwater-Home: *Alteromonadaceae*, Oceanospirillaceae, *Rhodobacteraceae*, and *Vibrionaceae* (Fig. 4). The enzymatic profile of each community followed similar trends, with the Marine-Home, Marine-Brackish and Freshwater-Brackish replicates having reduced enzyme activity (Suppl. Fig. 4) and more similar enzyme profiles relative to the Freshwater-Home enzyme profile (Fig. 2C).

*Biotic Filter:* In contrast to the extreme loss of abundant taxa caused by environmental filtering, the addition of interacting taxa from the two endmember communities into our Brackish media (‘Brackish-Coalescence’) had a more limited effect on microbial richness. Of the 967 Freshwater taxa found in the initial Freshwater-Home treatment, 37 were detected in the Brackish-Coalescence treatment. This set overlapped entirely with the set of taxa that survived through the environmental filter, with the exception of a single taxa that disappeared in the Freshwater-Brackish treatment but increased in abundance in response to the addition of an interacting community assemblage. There were 145 Freshwater-derived taxa that survived the Brackish treatment but did not persist when in the presence of the new interacting microbiome (lost in the Brackish-Coalescence treatment). Of the 588 Marine taxa detected in the original Marine-Home treatment, 171 were detected in the Brackish-Coalescence treatment. This set of Marine survivors overlapped considerably with the list of taxa that were tolerant of the environmental filter (with 143 taxa found in both taxa lists). There were 28 Marine-derived taxa who only persisted in Brackish media when also combined with the interacting microbiome (Brackish-Coalescence), and there were 302 Marine taxa that could survive the environmental filter (were found in Marine-Brackish treatments) but could not persist in the presence of the new blended microbiome (lost in the Brackish-Coalescence treatment).

We introduced at least 1528 taxa from both endmember inocula into the Brackish-Coalescence treatments (this is the sum of the distinct taxa derived from the two endmember communities). Given the rare biosphere constituents detected in our environmentally filtered treatments, we likely added a further 495 taxa, for a total taxa pool of >2000 microbial taxa. After coalescence, we detected only 472 taxa in the Brackish-Coalescence treatment. While richness declined roughly to the level of the Marine-Home, evenness was intermediate between the two endmembers, reflecting a shift in the shape of the dominance diversity curve. Taxa loss was not symmetric: only 37 of the original Freshwater microbiomes were detected, while 171 Marine-derived microbial taxa survived. Ten of these ‘surviving’ taxa were in common, and all ten of these overlapping taxa were members of the set of 21 taxa found in both original endmember inocula. One quarter (n=126) of the taxa detected in the Brackish-Coalescence treatment were not observed in the Marine-Brackish or Freshwater-Brackish treatments and thus we do not know from which endmember community they were derived. These rare biosphere constituents increased in abundance as a result of interactions between the endmember microbiomes.

The composition of the Brackish-Coalescence community overlapped almost entirely with the Marine endmember community (Fig. 2B). Both the Marine endmember, the Marine-Brackish and the Brackish-Coalescence communities were dominated by <u>*Alteromonas*</u>(~50% of relative abundance) (Fig. 3). Taxa that dominated the Freshwater microbiome were lost (Fig. 3, Fig. 5).). The enzymatic response followed a similar trend: the enzymatic profile for the Brackish-Coalescence microcosms was indistinguishable from the Marine-Brackish and slightly different from the Marine-Home enzyme profile, while significantly different from the enzymes of both the Freshwater-Home and Freshwater-Brackish treatments (Fig. 2C).

**Figure 5.**
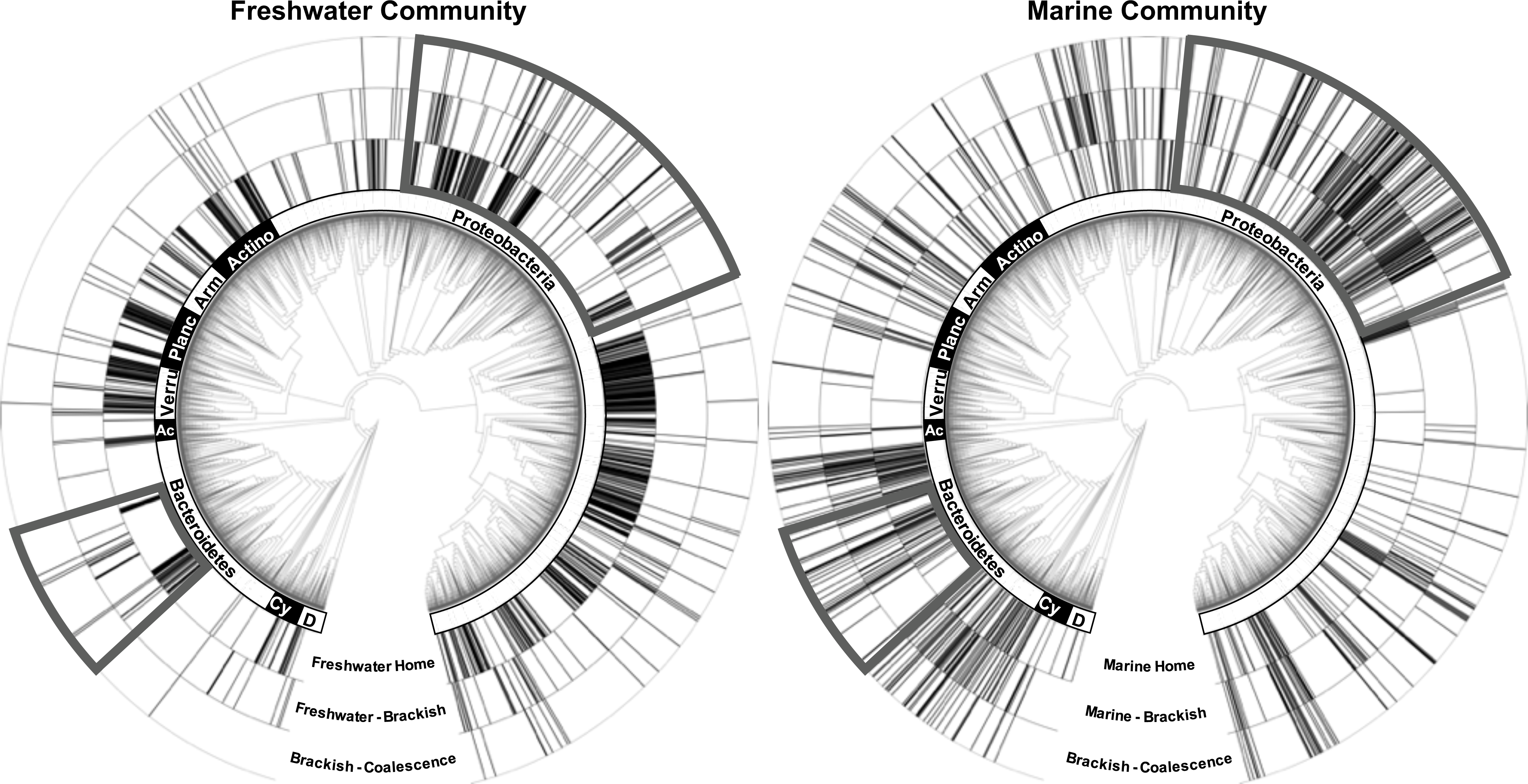
Community phylogenetic response to environmental filtering and to novel biotic interactions for Freshwater (left) and Marine (right) microbiomes. The <u>*inner ring*</u> shows the distribution of microbial taxa present in “home” conditions; <u>*middle ring*</u> represents the distribution of microbial taxa surviving the Brackish environment; and the <u>*outer ring*</u> displays the microorganisms of known origin surviving to Brackish-Coalescence. The gray boxes highlight phylogenetic regions where the Freshwater and Marine microbiomes exhibit distinct dispersion signals in response to the assembly filters. *The bacterial phyla are labeled inside the rings, with the following abbreviations ‘Actino’: Actinobacteria; ‘Arm’: Armatimonadetes; ‘Planc’: Planctomycetes; ‘Verru’: Verrucomicrobia; ‘Ac’: Acidobacteria; ‘Cy’: Cyanobacteria; ‘D’: Dependentiae*.

### Phylogenetic response to Coalescence varies by End Member Community

Taxa that were lost and gained from the Marine microbiome in response to environmental filter and to the biotic filter were closely related (Fig. 5). A sensitive taxon lost due to filtering was typically replaced by an increase in abundance of a tolerant sister taxa in the Marine microbiomes. In contrast, whole classes and orders of the Freshwater microbial community were lost and gained as a result of these two assembly filters. The phylofactorization of microbiomes exposed to the Brackish media identified nine factors, forming six non-overlapping clades capturing a total of 548 species (average 91 species) showed significant changes in the probability of being detected in Brackish water. All but one of these factors show an increase probability of being present relative to other Freshwater microorganisms (Suppl. Fig. 5). In contrast, there were four overlapping Marine clades (from five identified factors) totally 255 species (average 64 species) showing significant changes in detection probability with Brackish exposure (Suppl. Fig. 5).

## DISCUSSION

Environmental filters and biotic interactions were both important in determining which Freshwater and Marine microorganisms survived under Brackish common garden conditions. When transplanted from their home environments into Brackish media, both Freshwater and Marine communities lost >70% of all detectable taxa in the initial inoculum and home conditions. This was somewhat counter balanced by the emergence of the rare biosphere of each microbiome, accounting for 66% of the Marine microbiome and 82% of the Freshwater microbiome in the Brackish environment. The rare biosphere emergence stabilized Marine microbial richness to just over 100% of original richness. In contrast, Freshwater microbiome richness in Brackish conditions was only 21% of original richness, despite rare biosphere emergence. This awakening of the rare biosphere during Brackish exposure supports Paver *et al.* (2018), who show that certain rare microbial taxa that may possess wider salinity tolerance (‘crossing the salty divide’) may also be uniquely adapted to proliferating in these new environmental conditions. The biotic filter also imposed taxonomic shifts for both communities when added in combination to the Brackish arena. Under this Brackish-Coalescence treatment, a further 70.7% of the initial Freshwater inocula and 51% of the Marine inocula were not detected. Taken together, these patterns of major loss of specialists and counter balancing of emerging rare taxa explain the community convergence on the Marine microbiome.

There were significant differences in the taxonomic richness response of these two communities to our coalescence experiment. While both endmember communities were significantly different from their coalescent counterparts, only the Freshwater community had a significant reduction in taxa richness as a result of environmental filtering (Fig. 2C). This suggests that the Marine microbiome is, on the whole, more capable of dealing with the intermediate Brackish conditions for two main reasons: resistance of at least a third of the community to a range of salinity and nutrients, and resilience due to the high community buffering capacity of the Marine rare biosphere. The prevalence of Marine-derived rare Brackish tolerant taxa and consequential convergence towards the Marine microbiome is perhaps expected given the high physiological threshold of marine microbial taxa to a wide range of salinity (del Giorgio and Bouvier 2002, Wu *et al.* 2006, Herlemann *et al.* 2010). As we lose sensitive dominant taxa, we sample the rare biosphere more deeply. The rich ‘microbial seed bank’ buffers fluctuations in species richness (Lennon and Jones 2011, Jousset *et al.* 2017, Wang *et al.* 2017), which may help explain the limited functional change in this common garden experiment.

These two unique microbial communities show distinct phylogenetic responses to the environmental filters imposed by being transplanted into Brackish media. For our Freshwater community, whole clades turned over. We saw that several sensitive Freshwater clades were lost while multiple tolerant Freshwater clades become abundant enough to detect. In contrast, for our saltwater community the turnover was at a finer taxonomic resolution, with a loss and gain of sister taxa within clades. The composition of experimental replicates was remarkably similar, indicating that there are real differences in the ability of microbial taxa to survive transplant into altered salinities and that community composition responses are predictable. Community composition converged under Brackish conditions because for both endmember microbiomes there was a reservoir of tolerant rare taxa which increased in abundance when exposed to the intermediate environmental condition. The shift in composition towards the Marine microbiome resulted from a greater reservoir of tolerant taxa within that community. The evidence for this is both the compositional shift revealed through ordination and also the fact that the loss of initially detected taxa is not accompanied by a decline in species richness for the Marine microbiome.

From a fundamental science perspective, the immense contribution of the rare biosphere to the Brackish conditions is most fascinating and this rapid turnover of the community somewhat unique to microorganisms. With <2% of detectable overlap among the original microbiomes, the response to Brackish conditions converges the microbiomes to much higher taxonomic overlap, with the retention of the original overlapping taxa and immense emergence of the rare biosphere of each endmember community. This emergence of the microbial ‘seedbank’ and the dampened response to aggregate microbial function is an excellent example of microbial community resilience, where the rare biosphere plays a pivotal role in ecological rescue.

From an applied microbiology perspective and the natural history of microorganisms, learning which taxa are lost to environmental and biotic filtering will be instructive and useful for microbial engineering of ‘wild’ unculturable microbial taxa (Libby & Silver 2019). There are potential commercial applications for this experimental setup. The consistency of microbial community composition responses to experimental treatments suggests that this approach may prove quite useful in strategically identifying sensitive and tolerant taxa along many different real environmental gradients (Rocca *et al.* 2019). Over time, such information could lead to the development of microbial sensors, in which microbial community composition could be used to draw inferences about environmental conditions. By better understanding the resultant microbial community structure and function when microbial worlds collide, we may be able to better understand and modulate microbial communities in areas as wide as agricultural efficiency by microbial consortia (Busby et al 2017), bioremediation (Baez-Rogelio et al 2016, Sierocinski *et al.* 2017) or biomedical microbial transplants (Gibbons *et al.* 2017).

Our experiment also raises many new fundamental questions about the environmental and biotic processes that structure the microbiome. Because we observed compositional shifts at very fine levels of taxonomic resolution (~sister taxa) for our Marine microbiome when exposed to Brackish conditions, we speculate that biotic interactions are far more important within this community. Why this might be is an interesting question for microbial ecology.

## CONCLUSIONS

Our experimental approach to the study of microbial coalescence provides one of the first demonstration to directly and separately compare the relative strength of environmental and biotic filters in structuring intact microbial communities. In the case of mixing Fresh and Marine water, the Brackish intermediate condition proved to be a very strong environmental filter that had a greater effect on the Freshwater microbiome than its Marine counterpart. Applying this technique to other gradients may reveal cases in which biotic interactions dominate. Collectively, the use of this approach to detect the phylogenetic distribution of sensitivity and tolerance to various environmental gradients is likely to help us rapidly advance our ecological understanding of why microbial taxa live where they do.

## SUPPLEMENTARY INFORMATION

**Supplemental Figure 1.** Laboratory Experimental Incubation Setup. (Top) Reciprocal manipulation of communities into ‘home’, ‘away’ and ‘mixed’ environments, with and without community mixing. (Bottom) Field controls of: (Left) Intact communities with two levels of filtration to assess microbes-only and microbes + small predators; (Middle) Sterile environments of same filtration; (Right) Water-only negative controls. Replication is five microcosms per treatment.

**Supplemental Figure 2.** Nonmetric multidimensional scaling (NMDS) of bacterial community structure, demonstrating the minimal impacts of inoculum preparations and autoclaving environments on microbial community structure relative to unaltered positive control microcosms.

**Supplemental Figure 3.** 16S rRNA-based phylogeny of bacteria in the starting inocula. The inner ring shows SVs in the Freshwater inocula (green) and outer ring represents the Marine inocula (blue). Black arrows indicate the 21 overlapping SVs.

**Supplemental Figure 4.** Microbial function of Coalescing Microbial Communities vs. End member Controls. Barcharts of potential extracellular enzyme activity: (A) Aryl-sulfatase, (B) 1,4 Beta-glucosidase, (C) 1,4 N-acetylglucosidase, (D) Alkaline Phosphotase, (E) 1,4 Alphaglucosidase, (F) 1,4 Cellobiohydrolase, (G) D-xylosidase, and (H) L-leucine-aminopeptidase. Compact letter display represents pairwise comparisons among each of the five treatments, dotted lines denote the average of the end-point controls (Freshwater and Marine), and error bars display ±SE of each enzyme.

**Supplemental Figure 5.** Phylofactorization results for Brackish exposure on the Freshwater (Left) and Marine (Right) microbiomes. Top panels show the non-overlapping clades identified as significantly factors based on a Holm’s sequentially rejective 5% FWER cutoff, and bottom panels represent the corresponding phylogenetic location for each of these factors (6 in Freshwater, 4 in Marine).

