## Supplementary figures and images for "Rare Microbial Taxa Emerge When Communities Collide: Freshwater and Marine Microbiome Responses to Experimental Seawater Intrusion"

### Supplemental Figure 1

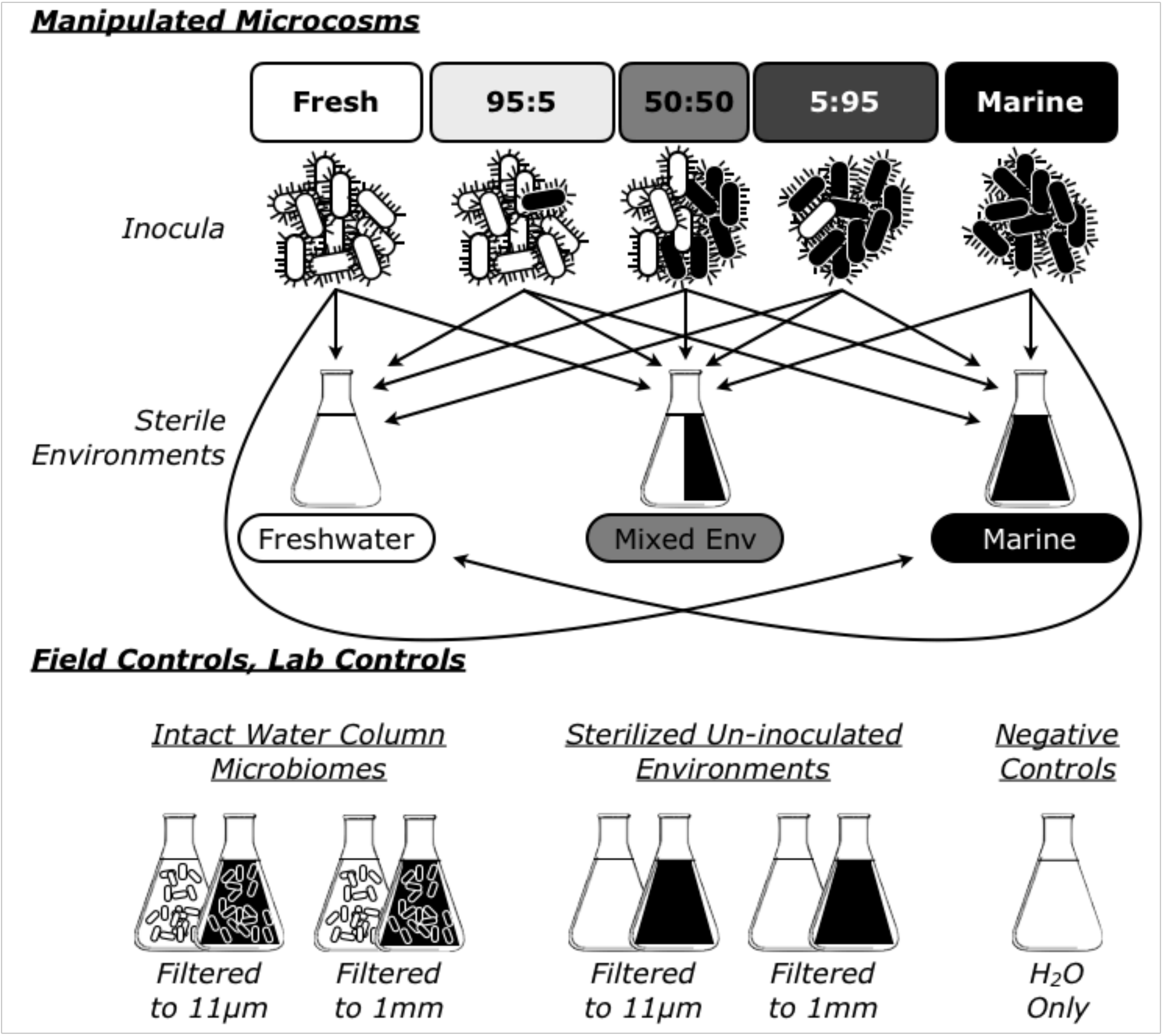

### Supplemental Figure 2

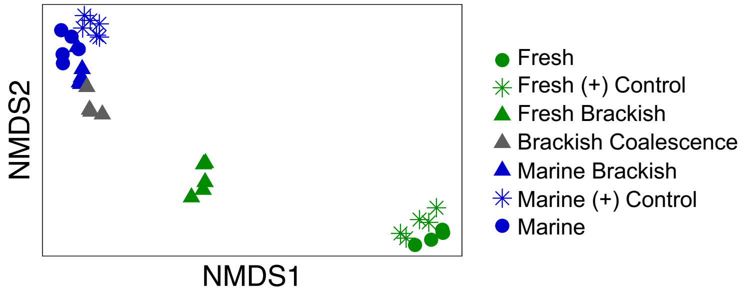

### Supplemental Figure 3

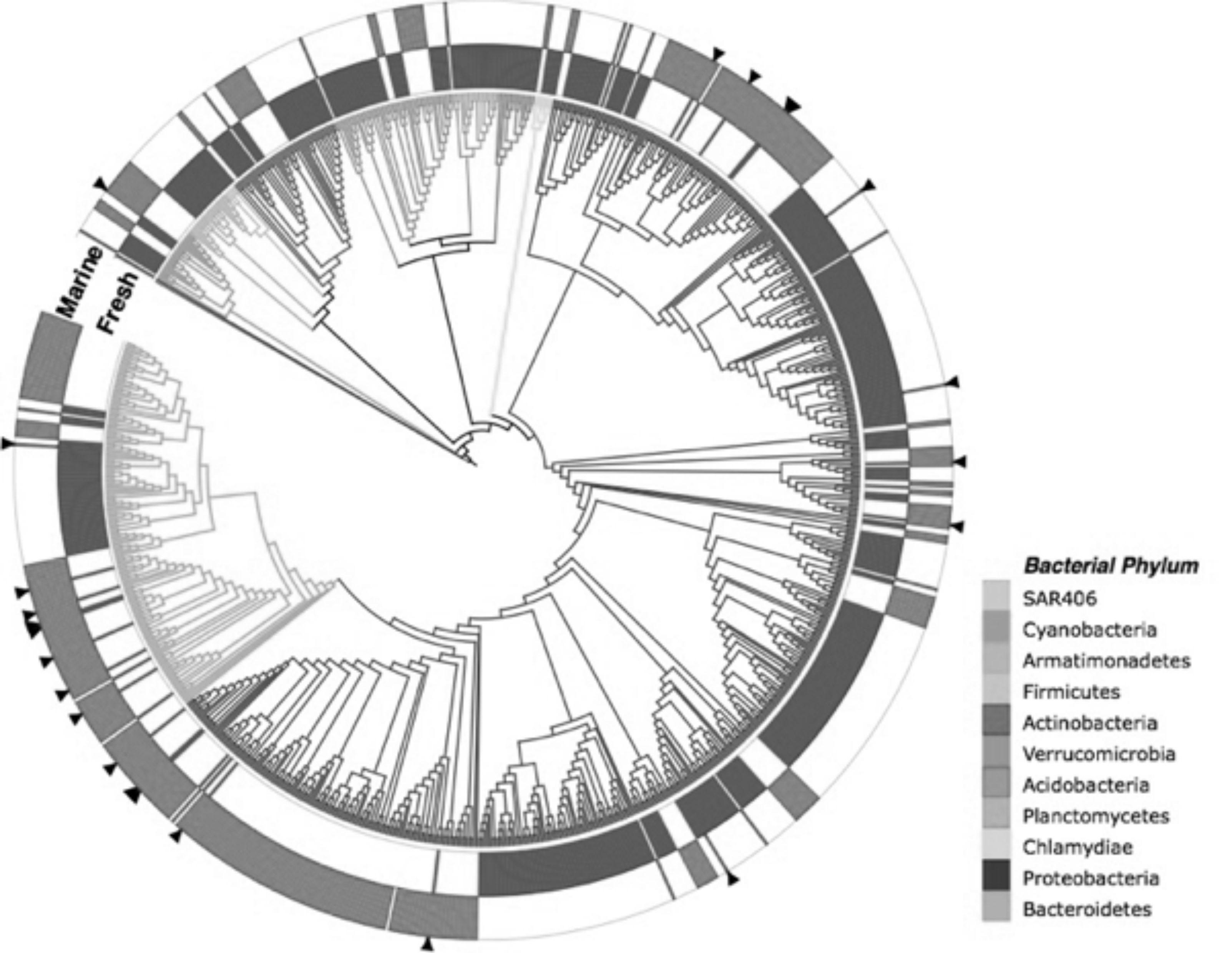

### Supplemental Figure 4

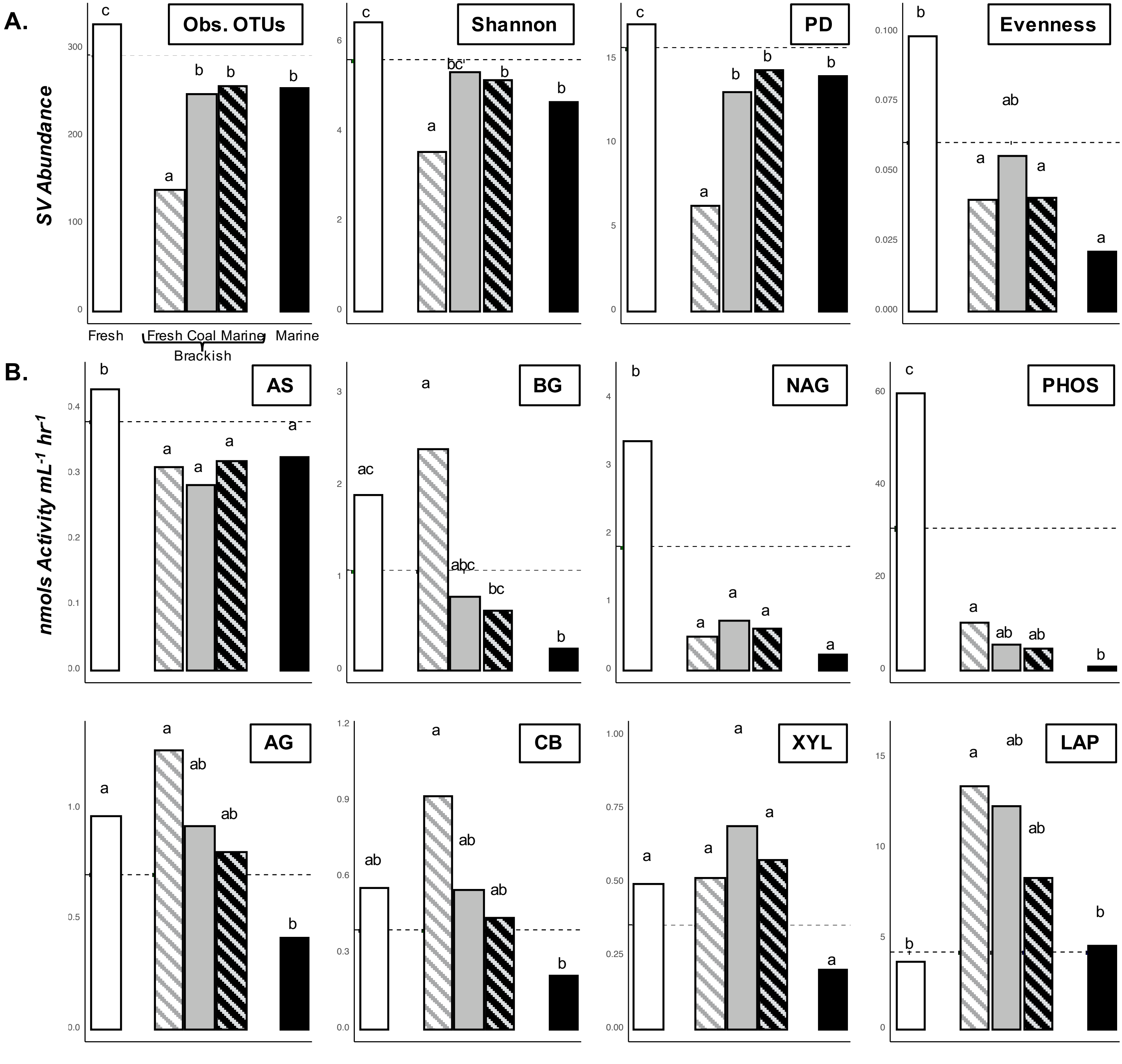

### Supplemental Figure 5

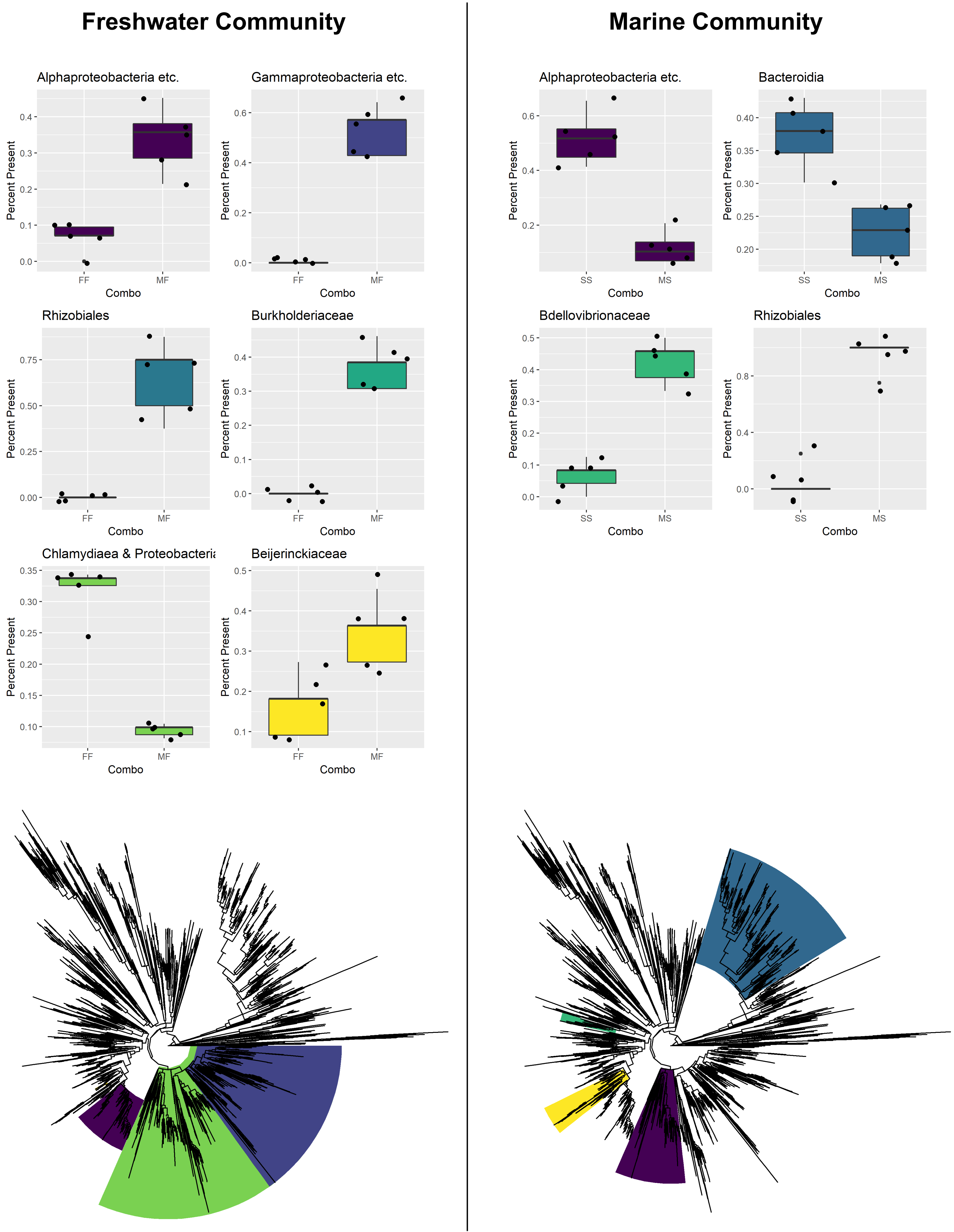
